# ROP16-mediated activation of STAT6 facilitates encystment of type III *Toxoplasma gondii* in neurons

**DOI:** 10.1101/2022.07.31.502204

**Authors:** Joshua A. Kochanowsky, Sambamurthy Chandrasekaran, Jacqueline R. Sanchez, Kaitlin K. Thomas, Anita A. Koshy

**Author notes:** Address correspondence to Anita Koshy,. Joshua A. Kochanowsky, University of California, Los Angeles, CA United States of America. Jacqueline R. Sanchez, Graduate Program in Neuroscience, University of Illinois Chicago.

## Abstract

*Toxoplasma gondii* establishes a long-lived latent infection in the central nervous system (CNS) of its hosts. Reactivation in immunocompromised individuals can lead to life threatening disease. Latent infection is driven by the ability of the parasite to convert from the acute-stage tachyzoite to the latent-stage bradyzoite which resides in long-lived intracellular cysts. While much work has focused on the parasitic factors that drive cyst development, the host factors that influence encystment are not well defined. Here we show that a polymorphic secreted parasite kinase (ROP16), that phosphorylates host cell proteins, mediates efficient encystment of *T. gondii* in stress-induced models of encystment and primary neuronal cell cultures (PNCs) in a strain-specific manner. Using short-hairpin RNA (shRNA) knockdowns in human foreskin fibroblasts (HFFs) and PNCs from transgenic mice, we determined that ROP16’s cyst enhancing abilities are mediated by phosphorylation of the host cell transcription factor STAT6. To test the role of STAT6 *in vivo*, we infected STAT6KO mice, finding that, compared to infected wild-type mice, infected STAT6KO mice have a decrease in cyst burden, but not overall parasite burden or dissemination to the CNS. Finally, we found a similar ROP16-dependent encystment defect in human pluripotent stem cell-derived neurons. Together, these findings identify a host cell factor (STAT6) that *T. gondii* manipulates in a strain-specific manner to generate a favorable encystment environment.

## Introduction

A broad range of microbes—viruses, bacteria, and parasites—establish long-term, latent infections by switching from a rapidly replicating state to a slow-growing quiescent one. While these chronic infections enable microbial spread to new hosts, they can also reactivate to cause overt disease when the host becomes immunocompromised. *Toxoplasma gondii,* a common obligate intracellular pathogen that infects most warm-blooded mammals including humans, switches from a rapidly replicating state, the tachyzoite, to a slow-growing, encysted state, the bradyzoite^1–5^. Bradyzoite-filled cysts are the hallmark of chronic infection and are primarily found in neurons in the central nervous system (CNS) and myocytes in skeletal and cardiac muscle^1,5^. For *T. gondii*, encystment allows infection of the definitive host (felids) and passage between intermediate hosts, thereby enabling both the sexual and asexual life-cycle of the parasite^5^. In humans, chronic infection is generally asymptomatic, but in the setting of acquired immune deficiencies, recrudescence can lead to severe pathology and even death. During the height of the HIV/AIDs epidemic, toxoplasmic encephalitis was the most common focal neurologic finding in AIDs patients^6,7^.

Given *T. gondii’s* clinical importance, many studies have focused on understanding persistence. These studies have identified many of the parasite factors that define and drive stage conversion and encystment and determined that a range of exogenous stresses (e.g., high pH) can trigger encystment in non-permissive cells (e.g., fibroblasts, macrophages)^8–11^. In permissive cells (neurons and myocytes), high levels of encystment can be achieved without the addition of exogenous stress^12–17^. Collectively, these data suggest that specific host cell pathways promote encystment, but only a single host gene (CDA-1) that influences encystment has been identified in myocytes^16,17^. Glutamine starvation and reactive oxygen species (ROS) production were recently identified as triggers of encystment in murine skeletal muscle cells and IFN-γ stimulated human induced pluripotent stem cells (iPSC)-derived glutamatergic neurons consistent with the observation that encystment may be a generalized response to stress^18,19^.

Though many *T. gondii* strain types encyst and cause clinically relevant disease^20–24^, most work on encystment has been done using type II strains. The ability to compare strains has highlighted that strain-specific effectors influence acute infection^25–29^, leading us to question whether strain-specific pathways that influence encystment might also exist. We were particularly interested in the polymorphic effector protein ROP16 for several reasons^30^. ROP16s from types I, II, and III translocate into the host cell nucleus and have a functional kinase domain, but only the type I and III alleles (ROP16_I_ and ROP16_III_ respectively), which are 99% identical in their amino acid sequence, cause prolonged phosphorylation and activation of the host transcription factors STAT3, STAT5a, and STAT6^30–34^. The importance of this STAT activation is strain-specific, as lacking ROP16 does not affect type I strain virulence in mice, but is essential for type III strain survival *in vivo*^31,35–36^. In addition, ROP16_III_-dependent phosphorylation of STAT6 dampens host cell ROS production in human and murine cells, enabling improved type III survival *in vitro*^37^. Considering the profound impact of ROP16 on host cell signaling and tachyzoite survival, we wondered if ROP16 might also influence encystment.

To address this possibility, we compared cyst formation in type II and type III strains that lacked ROP16 (IIΔ*rop16* and IIIΔ*rop16* respectively). We observed that deletion of ROP16 had no effect on type II encystment but significantly decreased type III encystment in both a stress model of encystment and in murine primary neuronal cell cultures (PNCs). Using IIIΔ*rop16* strains complemented with mutated ROP16s, we determined that the ROP16-dependent effect on type III encystment required a ROP16 that is capable of phosphorylating STATs. By using PNCs derived from mice deficient in STAT6 (STAT6KO)^38^ and infection of STAT6KO mice with wild-type type III parasites (WT_III_), we identified that efficient encystment required STAT6 *in vitro* and *in vivo*. Finally, we determined that ROP16 was also required for efficient encystment of type III parasites in human neurons. Together these results highlight a mechanism by which an allele of a parasite kinase (ROP16) facilitates encystment via activation of a host cell transcription factor (STAT6). To the best of our knowledge, this study identifies the first host cell gene that influences encystment in a strain-specific manner.

## Results

### ROP16 facilitates cyst development in a strain-specific manner

To assess how ROP16 affected encystment, we infected human foreskin fibroblasts (HFFs) with WT_III_, IIIΔ*rop16*, or strains where we re-introduced the type III allele of *rop16* (ROP16_III_) or the type II allele of *rop16* (ROP16_II_)^35–37^. We induced encystment by placing the infected cultures under alkaline stress (pH 8.2) in combination with CO_2_ deprivation (<5%) and tracked differentiation using immunofluorescent staining of *T. gondii* stage-specific surface antigens^39,40^ and *Dolichos biflorus* agglutinin (DBA), which binds sugar moieties present on the proteins that form the cyst wall^41^. At 6 days post infection (dpi) the WT_III_ strain no longer expressed the tachyzoite-specific antigen SAG1^39^, instead expressing the bradyzoite-specific surface antigen SRS9^40^, and stained positive for DBA indicating formation of the cyst wall (**Fig. S1**). In contrast, IIIΔ*rop16* parasites displayed a defect in SRS9 expression and DBA staining, which could be rescued by complementation of ROP16_III_, but not by complementation with ROP16_II_ (**Fig. S1**).

To further characterize and quantify this defect in cyst development, we optimized an automated imaging system using the Operetta^®^ CLS^TM^ platform and Harmony software (**Fig. 1A**). This automated system allowed us to track all parasitophorous vacuoles (PV) through staining with a polyclonal anti-*T. gondii* antibody^42^ and cysts through staining with DBA. Using this system, we tracked the encystment rate of our strains over the course of eight days. Compared to WT_III_ and ROP16_III_ parasites, IIIΔ*rop16* and ROP16_II_ parasites showed a 30-50% reduction in encystment at 4-8 dpi (**Fig. 1B**). To ensure that this decrease in encystment was not due to a decrease in overall parasite survival of the IIIΔ*rop16* and ROP16_II_ parasites, we used this automated system to track the accumulation of the total number of PVs over time. There were no significant differences in the accumulation of PVs between any of our strains, indicating that the defect in encystment we observed is unlikely to be explained simply by a defect in parasite survival in the mutant strains (**Fig. 1C**).

**Fig 1.**
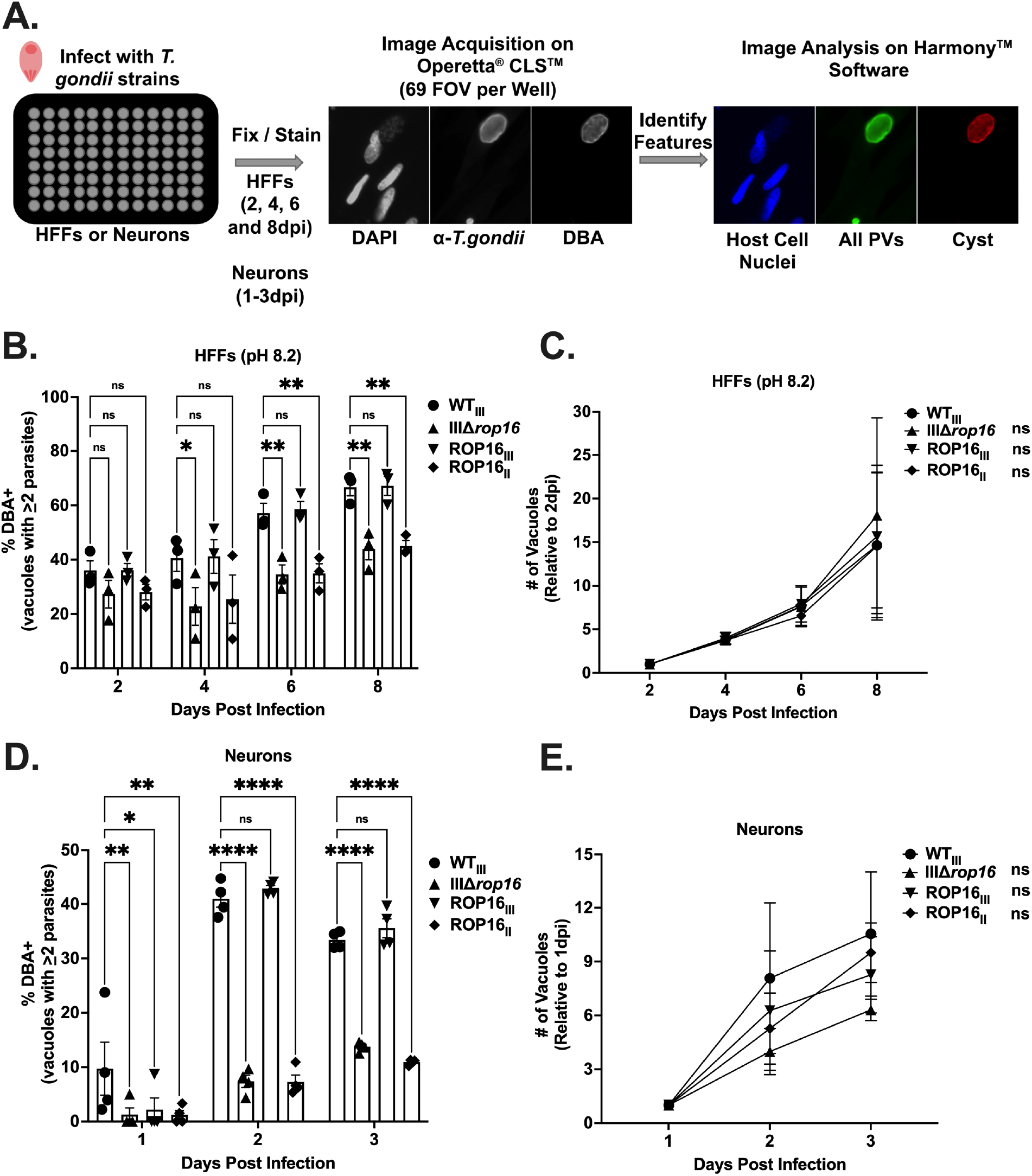
ROP16_III_ facilitates cyst formation of a type III parasites in multiple models of cyst development. (A) Schematic of automated cyst quantification using Operetta^®^ CLS™ and Harmony^®^ software. (B) Quantification of encystment over time in a stress model of encystment. % Cyst = (# of DBA^+^ Vacuoles with ≥2 parasites /# of Total Vacuoles with ≥2 parasites)*100. Bars, mean ± SEM. Black dots = Average % cyst for 1 experiment. (C) Quantification of accumulation of PVs over time in a stress model of encystment relative to 2 dpi. Dots, mean ± SEM. (D) Quantification of encystment over time in primary neuronal cultures (PNCs). Bars, mean ± SEM. Black dots = Average % cyst for 1 experiment. (E) Quantification of accumulation of PVs over time in PNCs relative to 1 dpi. Dots, mean ± SEM. (B, C) N = 10 wells/experiment, 3 experiments total. (D, E) N = 5 wells/experiment, 4 experiments total. (B-E) *p≤0.05, **p≤0.005, and ****p≤0.0001. ns = not significant, two-way ANOVA, Dunnett’s multiple comparisons test compared to WT_III_.

Although the alkaline and CO_2_ stress model is commonly used to increase encystment, it represents a highly artificial setting. As neurons are the major host cell type for persistent infection in the CNS^43–49^, we decided to use PNCs, which prior studies suggest induce encystment without the need for exogenous stress^12,14–15^. To test how ROP16 influenced type III encystment in neurons, we infected PNCs with WT_III_, IIIΔ*rop16*, or the two complemented strains and tracked cyst wall formation using DBA staining. Consistent with what we found in alkaline-stressed HFFs, we observed a defect in encystment in PNCs infected with IIIΔ*rop16* and ROP16_II_ compared to WT_III_ and ROP16_III_-infected cultures (**Fig. S2**). To quantify this defect more robustly, we used our automated cyst quantification system to track the encystment rate over 3 days. Compared to WT_III_ and ROP16_III_ parasites, IIIΔ*rop16* and ROP16_II_ parasites showed a 50-75% reduction in encystment from 1-3 dpi in PNCs (**Fig. 1D**). There were no significant differences in the accumulation of PVs between any of our strains in PNCs (**Fig. 1E**).

To determine if this ROP16-dependent defect in encystment was restricted to the type III strain or would also be seen in other *T. gondii* strain types, we used a ROP16 deficient type II strain (IIΔ*rop16*) generated using a CRISPR/CAS9 approach^35^. Unlike the defect observed in our IIIΔ*rop16*, deletion of ROP16 from a type II background did not result in a defect in cyst development in stressed HFFs or in PNCs (**Fig. S3A, B**). Together these results indicate that ROP16 is needed for efficient encystment of type III parasites, but not for type II parasites, suggesting that strain-specific differences in encystment exist.

### ROP16 facilitates encystment through host cell manipulations

While ROP16 is thought to primarily phosphorylate host cell proteins^30^, we sought to ensure that our cyst defect was not due to a direct effect of ROP16 on parasite proteins. To accomplish this goal, we tested whether encystment of the IIIΔ*rop16* strain could be restored by co-infection of a host cell with WT_III_ GFP-expressing parasites (WT_III_::GFP) (**Fig. 2A**). We hypothesized that if ROP16 facilitates cyst development through host cell manipulations then the IIIΔ*rop16* cyst defect should be rescued in host cells which also house WT_III_::GFP parasites that have injected a functional copy of ROP16. However, if ROP16 is required for phosphorylation of a parasite protein involved in encystment then co-infection with WT_III_::GFP parasites should be insufficient to restore encystment. Co-infection with WT_III_::GFP was sufficient to restore encystment of IIIΔ*rop16* in an alkaline stress model of encystment, indicating that ROP16 facilitates cyst development through host cell manipulations (**Fig. 2B**). Together these results indicate that ROP16 enhances encystment via host cell manipulations.

**Fig 2.**
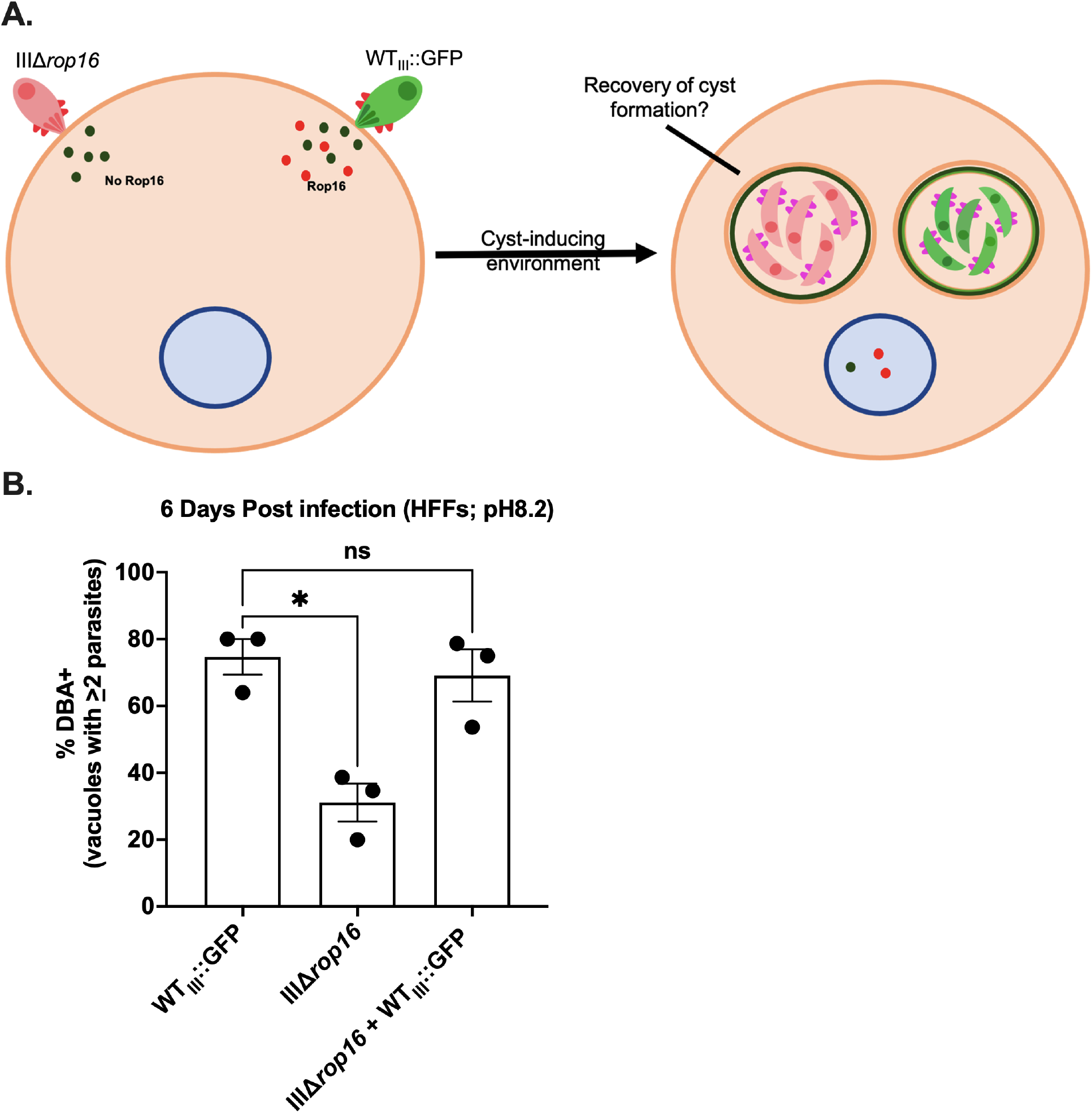
Co-infection of IIIΔ*rop16* with WT parasites restores encystment. (A) Schematic of co-infection experiment. (B) Quantification of encystment at 6 dpi in alkaline stress model of encystment. Bars, mean ± SEM. Black dots = Average % cyst for 1 experiment. N = 3 replicates/experiment, 3 experiments total. *p≤0.05, ns = not significant, one-way ANOVA, Dunnett’s multiple comparisons test compared to WT_III_.

### Efficient encystment requires a ROP16 with an active kinase domain and a leucine at position 503

Having determined that ROP16_III_-dependent host cell manipulation is essential for efficient type III encystment, we next sought to determine what domains or functions of ROP16 were required for this phenotype. To accomplish this goal, we used a previously generated panel of IIIΔ*rop16* complemented strains^37^. This panel of strains includes ROP16 mutants which are catalytically inactive (ROP16_IIIKD_) or lack a nuclear localization sequence (ROP16_IIIΔNLS_) (**Fig. 3A**)^37^. In addition, as prior work demonstrated that a single polymorphic amino acid on ROP16 determined the strain-specific activation of STAT3/5a/6^34^, this panel also included strains which express a ROP16_III_ in which the leucine at position 503 was switched to a serine, rendering it “STAT-dead” (ROP16_IIISD_), and a strain that expresses a ROP16_II_ in which the serine at position 503 was changed to a leucine, rendering it “STAT-active” (ROP16_IISA_)^37^ (**Fig. 3A**). We tested the ability of this panel of parasites to encyst in the stress model of encystment and in PNCs. In both models, IIIΔ*rop16*, ROP16_II_, ROP16_IIISD,_ and ROP16_IIIKD_ parasites were defective in forming cysts compared to WT_III_, ROP16_III_, or ROP16_IISA_ parasites (**Fig. 3B, C**). Together these data indicate that efficient *in vitro* encystment is dependent on kinase activity and a leucine at position 503, while the nuclear localization sequence (NLS) is dispensable. Additionally, the ability of ROP16_IISA_ to restore these defects suggested a role for activation of STATs in cyst development.

**Fig. 3.**
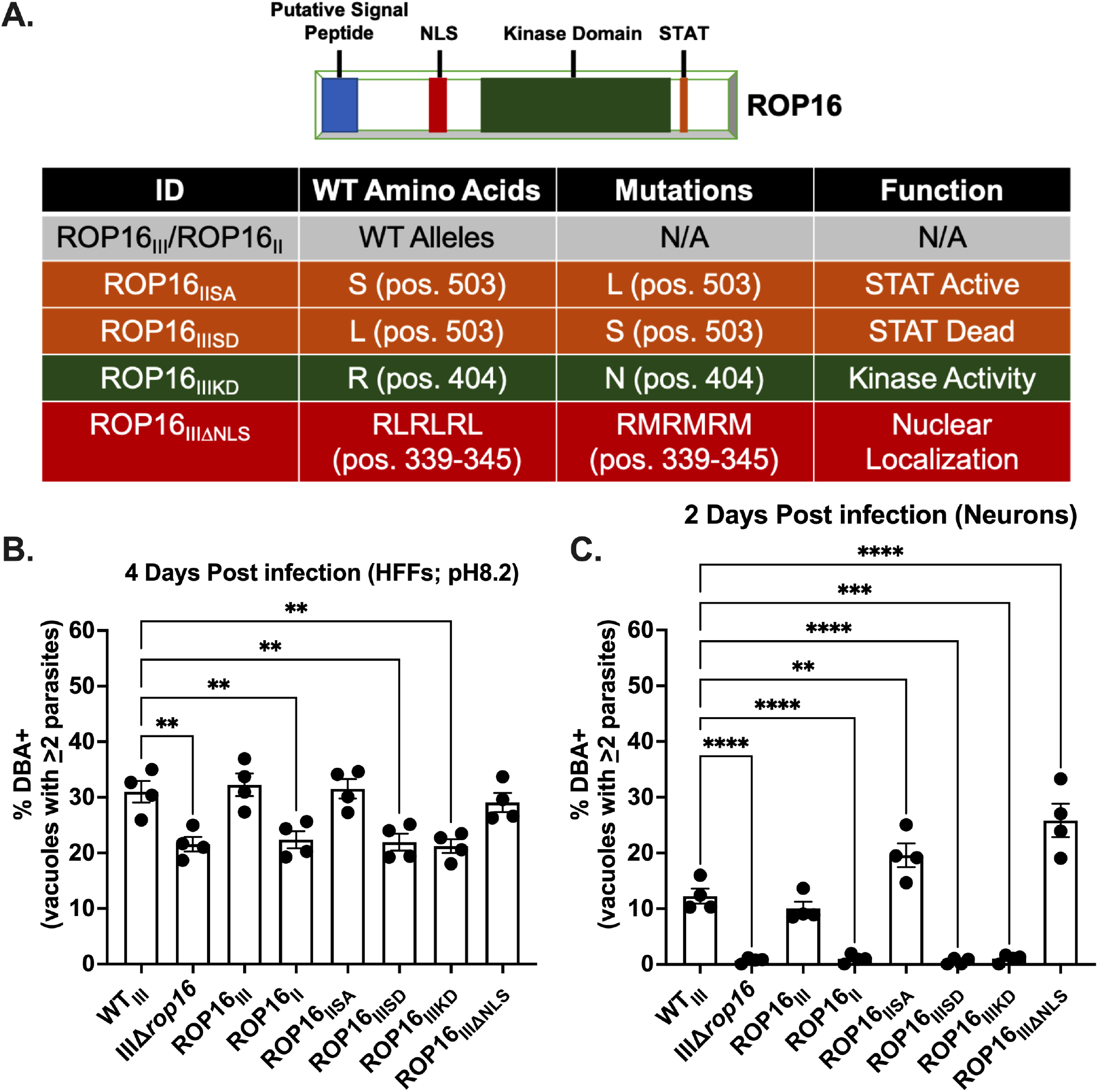
Efficient encystment requires a ROP16 with a functional kinase domain and a leucine at position 503. (A) Schematic of ROP16 and mutations in ROP16 constructs. (B) Quantification of encystment at 4 dpi in alkaline stress model of encystment. Bars, mean ± SEM. Black dots = Average % cyst for 1 experiment. N = 10 wells/experiment, 4 experiments total. (C) Quantification of encystment at 2 dpi in PNCs. Bars, mean ± SEM. Black dots = Average % cyst for 1 experiment. N = 3 wells/experiment, 4 experiments total. (B), (C) **p≤0.005, ***p≤0.0005, and ****p≤0.0001, one-way ANOVA, Dunnett’s multiple comparisons test compared to WT_III_.

### STAT6 is required for efficient encystment *in vitro*

To test the role of STATs in type III encystment, we used previously generated STAT3, 5a, and 6 knock-down HFFs^37^ in a cyst assay. Lentiviral-mediated knock-down of STAT6 significantly reduced encystment of WT_III_ parasites at both 2 and 4 dpi compared to parasites infecting non-targeting shRNA control HFFs. (**Fig. 4A**). Knockdown of STAT5a reduced encystment only at 4 dpi, while knock-down of STAT3 had no effect on encystment at 2 or 4 dpi (**Fig. S4**).

**Fig. 4.**
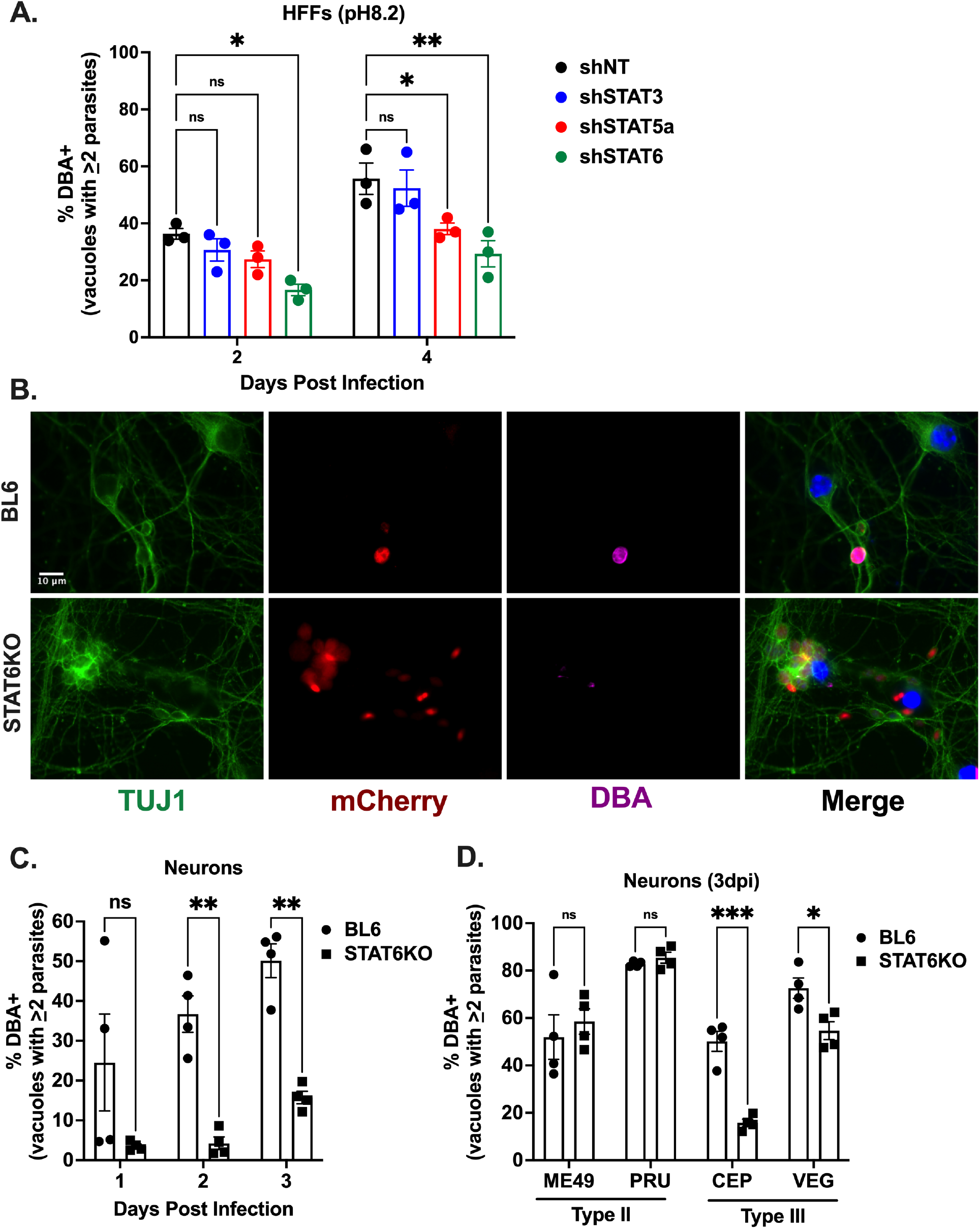
STAT6 facilitates encystment of type III, but not type II parasites *in vitro*. (A) Quantification of encystment at 2 and 4 dpi in alkaline stress model of encystment. Bars, mean ± SEM. N = 3 replicates/experiment, 3 experiments total. (B) IFA of cyst assay. BL6 (control) or STAT6KO neurons were infected with WT_III_ (MOI 0.1) for 2 dpi. Images depict anti-TUJ1 (green), mCherry (red), DBA (magenta), and DAPI (blue). (C) Quantification of encystment of WT_III_ parasites overtime in PNCs. Bars, mean ± SEM. N = 5 wells/experiment, 4 experiments total. (D) Quantification of encystment at 3 dpi of multiple WT_II_ and WT_III_ parasites in PNCs. Bars, mean ± SEM. N = 5 wells/experiment, 4 experiments total. (A) *p≤0.05 and **p≤0.005. ns = not significant, two-way ANOVA, Dunnett’s multiple comparisons test compared to shRNA non-targeting control (shNT). (C) **p≤0.005. ns = not significant, two-way ANOVA, Dunnett’s multiple comparisons test compared to BL6. (D) *p≤0.05 and ***p≤0.0005. ns = not significant, one-way ANOVA, Dunnett’s multiple comparisons test compared to BL6.

As the STAT6 knock-down showed the strongest effect on cyst formation and STAT6KO mice are commercially available, we decided to test the role of STAT6 in cyst development using PNCs from STAT6KO mice^38^. We observed a significant decrease in encystment of WT_III_ parasites in STAT6KO PNCs at all time points compared to parasites in PNCs from BL6 control mice (**Fig. 4B, C**). To determine if host cell STAT6 was required for efficient encystment in other *T. gondii* strains, we infected STAT6KO PNCs with two type II strains (ME49 and Prugniaud (PRU)) and two type III strains (CEP and VEG). At 3 dpi neither of the type II strains showed a defect in encystment, while both type III strains displayed a statistically significant decrease in encystment in STAT6KO PNCs (**Fig. 4D**). Taken together these results indicate that STAT6 is required for maximal cyst development in HFFs and PNCs for type III strains, but not type II strains.

### STAT6 is required for efficient encystment of type III parasites *in vivo*

As previously noted, deletion of ROP16 from type III strains causes the parasite to be cleared during acute infection which results in a significant decrease in parasite dissemination to the CNS and thus cyst burden^35,36^, making it difficult to assess the role of ROP16 during chronic infection *in vivo*. However, the identification of STAT6 activation as the downstream mediator of the ROP16 encystment phenotype *in vitro* opened an avenue for assessing the role of STAT6 activation during chronic infection *in vivo*. To test the role of STAT6 in chronic infection, we infected STAT6KO mice and BL6 control mice with type III (CEP) parasites and harvested brains at 17 dpi to measure parasite and cyst burden (**Fig. 5A**). When we quantified parasite burden by quantitative PCR (qPCR) for the *T. gondii*-specific gene B1, we observed no significant difference between STAT6KO and BL6 control mice (**Fig. 5B**). In contrast when we quantified cyst burden via immunofluorescent staining of infected CNS tissue, we observed a significant decrease in CNS cyst burden in STAT6KO infected mice compared to BL6 controls (**Fig. 5C**).

**Fig. 5.**
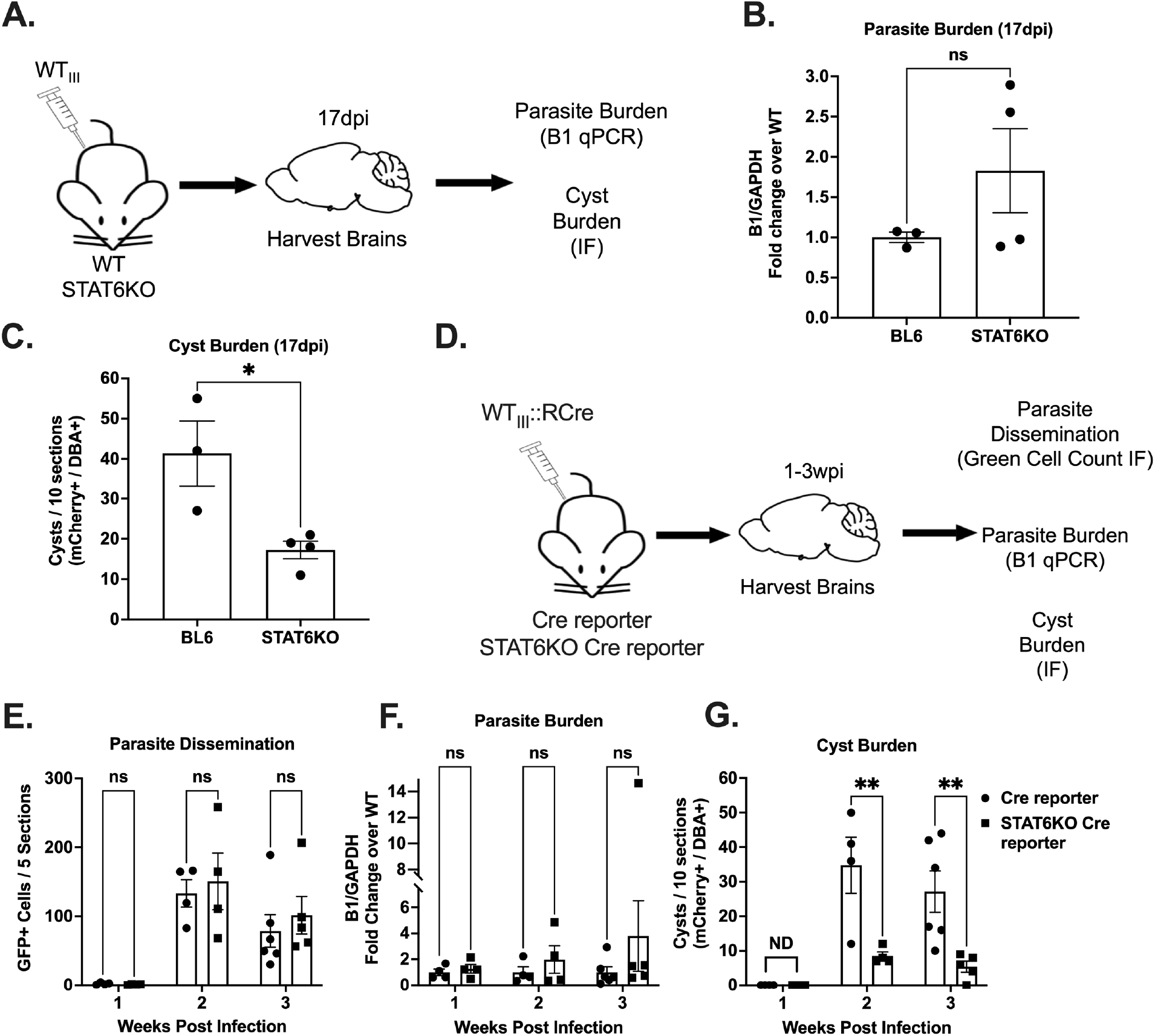
STAT6KO mice have lower cyst burdens despite having similar parasite dissemination and burden in the CNS. (A) Schematic of mouse infection for STAT6KO mice. (B) Quantification of *T. gondii* gene B1 was done using qPCR. Bars, mean ± SEM. Black dots = 1 mouse. N = 3-4 mice per condition. (C) Quantification of cyst burden. Total number of cysts per 10 sections. Bars, mean ± SEM. Black dots = 1 mouse. N = 3-4 mice per condition. (D) Schematic of mouse infection for Cre reporter (control) and STAT6KO Cre reporter mice. (E) Quantification of the total number of GFP^+^ cells per 5 sections per mouse. Bars, mean ± SEM. Black dots = 1 mouse. N = 4-6 mice per condition per timepoint. (F) Quantification of *T. gondii* gene B1 as in (B). Bars, mean ± SEM. Black dots = 1 mouse. N = 4-6 mice per condition per timepoint. (G) Quantification of cyst burden as in (C). Bars, mean ± SEM. Black dots = 1 mouse. N = 4-6 mice per condition per timepoint. (B, C) *p≤0.05. ns = not significant, one-way ANOVA, Dunnett’s multiple comparisons test compared to BL6. (E-G) **p≤0.005. ns = not significant, two-way ANOVA, Dunnett’s multiple comparisons test compared to Cre-reporter.

We next sought to determine if our cyst defect was secondary to differences in parasite dissemination to the CNS. To address this possibility, we generated STAT6KO Cre reporter mice by crossing STAT6KO mice to Cre reporter mice^50^ that express a green fluorescent protein (GFP) only after the cells have undergone Cre-mediated recombination (**Fig. S4**). We infected control Cre reporter mice and STAT6KO Cre reporter mice with parasites that express Cre recombinase fused to a parasite protein (WT_III_::RCre)^49,51^ that is injected into the host cell before invasion. As this system allows us to mark and track cells injected with *T. gondii* protein, the quantification of GFP^+^ CNS cells allows us to track parasite dissemination to the CNS. At 1-3 weeks post infection (wpi), we harvested brains to assess parasite dissemination, CNS parasite burden, and CNS cyst burden (**Fig. 5D**). There was no significant difference in the CNS GFP^+^ cell count at any timepoint between STAT6KO Cre reporter and control Cre Reporter mice (**Fig. 5E**). Consistent with the prior cohort (**Fig. 5B**), we observed no significant difference in CNS parasite burden between STAT6KO Cre reporter and control Cre reporter mice at any timepoint (**Fig. 5F**). In contrast to the GFP^+^ cell count and qPCR data, we observed a significant decrease in CNS cyst burden at 2 and 3 wpi in mice that lacked STAT6 (**Fig. 5G**). Together these results show that STAT6 facilitates encystment *in vivo* and leads to a reduction in cyst burden during chronic infection without affecting dissemination of the parasite or overall parasite burden in the CNS.

### ROP16 facilitates encystment in human neurons

Having shown that ROP16 facilitates encystment in murine PNCs and in the CNS *in vivo*, we wanted to know if ROP16 also affected cyst development in human neurons. To address this possibility, we differentiated human pluripotent stem cells into neurons (hPSC neurons) and then infected them with WT_III_ or IIIΔ*rop16* parasites for 1-3 days. We observed a decrease in cyst formation by immunofluorescent staining at 3 dpi in hPSC neurons infected with IIIΔ*rop16* compared to WT_III_ parasites (**Fig. 6A**). When we quantified encystment over time, we saw a significant defect in encystment (∼66% reduction) at 2 and 3 dpi in the IIIΔ*rop16* compared to WT_III_ parasites (**Fig. 6B**). These results suggest that, like murine neurons, ROP16 facilitates encystment of type III parasites in human neurons.

**Fig. 6.**
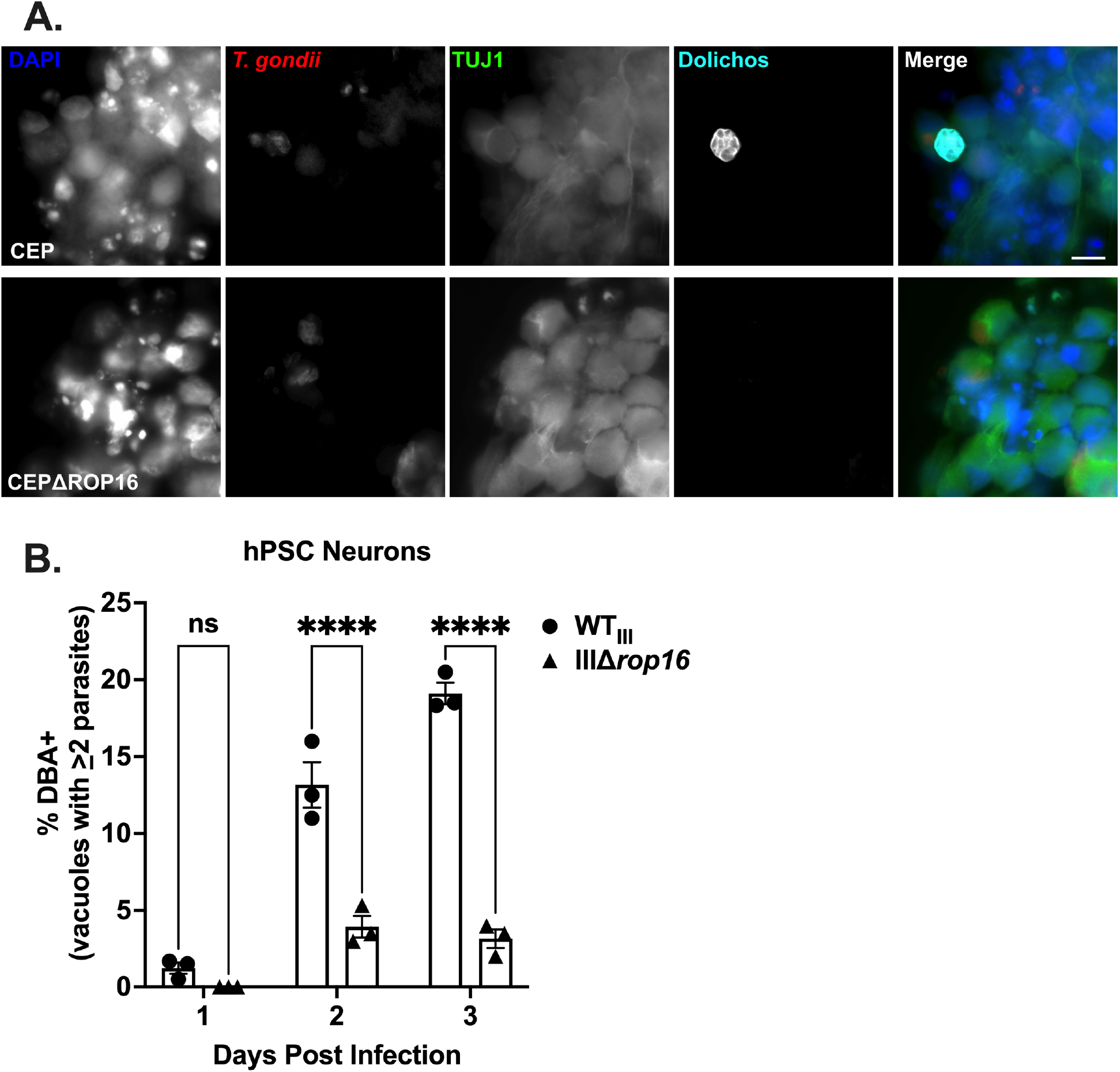
Deletion of ROP16 impairs cyst development in human neurons. (A) IFA of infected (3 dpi) hPSC neurons. Images depict anti-TUJ1 (green), mCherry (red), DBA (cyan), and DAPI (blue). (B) Quantification of encystment at 1-3 dpi in human stem cell derived neurons. Bars, mean ± SEM. N = 3 replicates/experiment, 3 independent experiments. **p≤0.005. ns = not significant, two-way ANOVA, Dunnett’s multiple comparisons test compared to WT_III_. Scale bar = 10µm.

## Discussion

The data presented here show that ROP16 mediates efficient encystment of type III parasites via activation of STAT6 in human and murine *in vitro* models of encystment and in mice *in vivo*. STAT6 is the second host gene to be identified as playing a role in encystment and the first to show a strain-specific effect on encystment. The importance of this latter point is that it questions the assumption that encystment is a process largely conserved between *T. gondii* strains.

This work emphasizes that parasite-host cell outcomes often vary by cell type and context. In non-neuronal cells, ROP16-triggered activation of STAT6 has been shown to influence tachyzoite survival by suppressing IFN-γ-induced nitric oxide production and IFN-γ-independent ROS production^31,37^. The data presented here show that in conditions that favor encystment (HFFs grown under alkaline stress and neurons), activation of STAT6 influences parasite stage conversion. These ROP16-mediated effects are presumptively driven by STAT6 altering gene expression in a cell-specific and context-dependent manner. Future studies will focus on defining the STAT6-dependent neuronal genes that promote encystment of type III strains.

The identification of a role for ROP16 in cyst development also raises the question of what other *T. gondii* secreted effector proteins prime cells to facilitate encystment. Until now, *T. gondii’s* repertoire of effector proteins have largely been studied in tachyzoite conditions and during acute infection. Much less is known about how *T. gondii*’s manipulation of host cells influence other stages of the parasite’s lifecycle. As ROPs are largely thought to be secreted only during the initial invasion process^52^, any effects on cyst development are likely to be involved in priming the host cell to influence the early events of stage conversion. The dense granule proteins (GRAs) are known to be secreted post invasion^53^, with evidence suggesting that at least one of these GRAs (TgIST) is secreted post cyst formation^54^. Thus, these effectors may play a larger role in cyst development and maintenance. As seen in this work, understanding how *T. gondii* effectors influence cyst development may provide a mechanism for identifying more host cell pathways and proteins that enable encystment.

Finally, ROP16 appears to be dispensable for encystment of type II parasites, indicating that there are strain-specific differences in stage conversion and cyst development. Prior strain-specific differences in stage conversion have been observed and were linked to parasite replication rates^55,56^. The data presented here suggest an additional level of complexity—different *T. gondii* strain types may require strain-specific host cell manipulations to produce optimal bradyzoite-inducing conditions. Given that type II strains lack a ROP16 that activates STATs and thus do not cause prolonged STAT6 activation^30,34^ this raises the question: do type II strains have unique host cell requirements for encystment? Or have they developed alternative mechanisms to activate the same downstream, encystment-promoting host genes? The availability F1 progeny from type II x type III cross^30^ in combination with our newly developed automated cyst imaging protocol opens a way to use quantitative trait loci mapping as a powerful tool to identify more parasite genes and associated host pathways that influence strain-specific differences in encystment.

In summary, the data presented here show that ROP16-dependent activation of STAT6 mediates efficient encystment of *T. gondii* in a strain-specific manner. These findings highlight that the outcome of host cell manipulation varies by host cell type and context. They also highlight that there are likely to be multiple layers of control for achieving maximal encystment and that these layers may vary between *T. gondii* strains.

## Methods

### Parasite Maintenance

As previously described, all parasite strains used in this study were maintained through serial passage in human foreskin fibroblasts (gift of John Boothroyd, Stanford University, Stanford, CA) using DMEM, supplemented with 10% fetal bovine serum, 2 mM glutagro, and 100 IU/ml penicillin and 100 μg/ml streptomycin. In addition to the strains described below, the previously described type III (CEP), IIIΔ*rop16* strains, and ROP16_III_ parasites expressing mCherry or tdTomato were used^35–37,49^.

### Generation of ROP16 Mutant Strains

Generation of IIIΔ*rop16* and ROP16 mutant parasites were previously described^37^. For the IIΔ*rop16* strain, type II parasites (PRU) were co-transfected the sgROP16Up CRISPR and sgROP16Down CRISPR plasmids along with the pTKO plasmid containing ROP16 homology regions surrounding a hypoxanthine phosphoribosyltransferase (*hxgprt)* cassette^35^. Parasites were then screened for expression of *hxgprt* using media containing 25 mg/ml mycophenolic acid and 50 mg/ml xanthine, prior to cloning by limiting dilution^57^.

### Mice

All procedures and experiments were carried out in accordance with the Public Health Service Policy on Human Care and Use of Laboratory Animals and approved by the University of Arizona’s Institutional Animal Care and Use Committee (#12-391). All mice were bred and housed in specific-pathogen-free University of Arizona Animal Care facilities. Cre reporter mice (stock no. #007906) and STAT6KO mice (stock no. 005977) were originally purchased from Jackson Laboratories. STAT6KO Cre reporter homozygotes were generated by breeding STAT6KO and Cre reporter mice and resulting heterozygotes progeny. The homozygosity at both loci were confirmed by PCR (**Fig S4**). Mice were inoculated with 10,000 freshly syringe-lysed parasites diluted in 200 μl of UPS grade PBS. Age- and sex-matched mice were used.

### Primary murine neuron culturing

Mouse primary cortical neurons were harvested from E17 mouse embryos obtained from pregnant Cre reporter mice or STAT6KO. Dissections of E17 cortical neurons were performed as described previously. Primary neuronal cell cultures were generated by methods described previously with minor modifications^58^. The culture plates were prepared by coating with 0.001% poly-L-lysine solution (Millipore Sigma, P4707, diluted in water 1:10) for plastic surfaces and 100 µg/ml poly-L-lysine hydrobromide (Sigma, P6282, dissolved in borate buffer, pH 8.4) for glass surfaces overnight. They were washed three times for 10 minutes each with water and transferred to plating media (MEM, 0.6% D-glucose, 10% FBS). Neurons were seeded at appropriate densities: 100,000 in 24 well plates with coverslips for confocal imaging and 20,000 in 96 well plates for use on the Operetta platform. Four hours after plating, full media exchange to neurobasal media (Neurobasal base media, 2% B27 supplement, 1% L-glutamine and 1% penicillin-streptomycin) was performed. On day in vitro (DIV) 4, neurons received a half volume media change of neurobasal media with 5 µM cytosine arabinoside (AraC, Millipore Sigma, C6645) to stop glial proliferation. One third media exchanges with neurobasal media occurred every 3-4 days thereafter. All the experiments were performed on 10 DIV neurons.

### Cyst Assays

Confluent HFF monolayers and primary neurons were cultured on glass coverslips, 96 well plates (Perkin Elmer; 6055302), or 6 well plates as indicated and infected with parasites at the MOIs indicated in the figure legends. For the alkaline stress model^10^, HFFs were infected with parasites for 4 hours under non-cyst inducing conditions (pH 7.1 and CO_2_ = 5%) to allow the parasites to invade. After 4 hours, the media was changed to cyst inducing conditions (50 mM HEPES [pH 8.1] in RPMI supplemented with 1% fetal bovine serum, penicillin, and streptomycin). Media changes occurred every 2 days to maintain pH levels. For primary neuronal cell cultures, parasites were simply added to culture at the indicated MOIs. For Operetta^®^ analyses, entire wells were imaged at 20x (69 total fields of view). 5-10 wells were imaged per experiment depending on the assay. Cells were fixed at the indicated timepoints and stained as described below. Images were then analyzed with Harmony™ software. Briefly, host cell nuclei were identified and enumerated via DAPI staining and the standard nuclei identification module. PVs were stained using an antibody that stains the PVM (Invitrogen PA17252). Stained PVs were identified using the Harmony™ find spots tool. Fluorescent intensity cutoffs were set at 10,000 and length and width cutoffs were set at 5 μm and 2 μm respectively. For purposes of cyst quantification, we wanted to exclude vacuoles with fewer than 2 parasites as these could be recent infections that have not yet had time to form cysts. To identify vacuoles with ≥2 parasites length and width cutoffs were set at 8 μm and 4 μm respectively. To identify cysts, we used the identify image area tool on DBA^+^ PVs identified as having ≥2 parasites. The intensity cutoff for positive cyst staining was set at 7,500 as determined by taking the average DBA background signal for parasites stained under non-cyst inducing conditions. The percent encystment in each well was then calculated by taking the number of DBA^+^ PVs (≥2 parasites) divided by the total number of PVs (≥2 parasites) followed by multiplication by 100. All Harmony™ Analysis programs are available upon request.

### Immunofluorescence Microscopy

For immunofluorescence assays, cells were fixed with 4% paraformaldehyde for 15 min. The cells were then permeabilized and blocked for 60 min with 0.1–0.2% (vol/vol) Triton-X 100 in phosphate buffered saline (PBS) (pH 7.4), typically with 3% (wt/vol) goat serum. For the alkaline-stress induced model of encystment, cells were incubated overnight with rabbit anti-*T. gondii* polyclonal antibody (Invitrogen PA17252) at 1:10,000; or mouse anti-SAG1 (DG52)^39^ at 1:10,000 and rabbit anti-SRS9^59^ (gifts of John Boothroyd, Stanford University, Stanford, CA) at 1:10,000; and Biotinylated-DBA at 1:500, followed by incubation with Strepavadin-647, Alexa Fluor-488, and Alexa Fluor-568 secondary antibodies at 1:500 (Molecular Probes) for 1 hour. Coverslips were mounted on slides using Fluoromount-G (ThermoFisher, 00-4958-02). For primary neuronal culture, cells were stained with mouse anti-TUJ1 (Millipore Sigma, MAB1637) at 1:500, rabbit anti-*T. gondii* polyclonal antibody (Invitrogen PA17252) at 1:10,000 and Biotinylated-DBA at 1:500, followed by incubation with Strepavadin-647, Alexa Fluor-488, and Alexa Fluor-568 secondary antibodies at 1:500 (Molecular Probes) for 1 hour. For 96 well plates wells were covered with 100μLs of Fluoromount-G diluted 1:50 in 1x PBS.

### Tissue preparation for histology and DNA extraction

At appropriate times post infection, mice were anesthetized with ketamine (24 mg/ml) and xylazine (4.8 mg/ml) cocktail and transcardially perfused with ice cold PBS. After harvesting organs, the left half of the mouse brain was fixed in 4% paraformaldehyde in phosphate buffer and kept at 4°C overnight, rinsed in PBS, and then was embedded in 30% sucrose. Post fixation, brains were sectioned into 40 μm thick sagittal sections using a freezing sliding microtome (Microm HM 430). Sections were stored in the cryoprotective media (0.05 M sodium phosphate buffer containing 30% glycerol and 30% ethylene glycol) as free-floating sections until stained and mounted on to slides. The right half of the brain was sectioned coronally into 2 halves and stored in separate tubes. These tubes were flash frozen and stored at −80°C until used for DNA extraction.

### Quantitative real time PCR

For quantification of parasite burden, genomic DNA from the rostral quarter of the frozen brain was isolated using DNeasy Blood and Tissue kit (69504, Qiagen) and following the manufacture’s protocol. The *T.gondii*-specific, highly conserved 35-fold repeat B1 gene was amplified using SYBR Green fluorescence detection with the Eppendorf Mastercycler ep realplex 2.2 system. GAPDH was used as control to normalize DNA levels. Results were calculated as previously described^35^.

### Green Cell and Cyst Counts

For enumeration of CNS cells that had undergone Cre-mediated recombination, sagittal brain sections were washed and mounted as previously described^35^. The total number of GFP^+^ cells were enumerated by using a standard epifluorescent microscope (EVOS microscope). Analyses were performed on 3 brain sections per mouse, after which the resulting numbers were then averaged to obtain the mean number of GFP^+^ cells/section/mouse. For enumeration of cysts, sagittal brain sections were washed and blocked in 3% Goat Serum in 0.3% TritonX-100/TBS for 1hour. Sections were then incubated with biotinylated DBA (Vector Laboratories 1031, 1:500) for 24 hours, followed by incubation with 647 Streptavidin (Invitrogen, 1:2,000) for 1 hour. Sections were mounted as described above. The number of cysts were enumerated using a standard epifluorescent microscope (EVOS microscope). Only objects that expressed mCherry/tdTomato and stained for DBA were quantified as cysts. Investigators quantifying were blinded to the infection status of the mouse until after the data were collected.

### Generation of hPSC Neurons

H7 human embryonic neural stem cells (NSCs) derived from the NIH-approved H7 embyronic pluripotent stem cells (WiCell WA07) were purchased from the University of Arizona iPSC core (https://stemcells.arizona.edu/) and expanded on NSC expansion medium (NEM) (Thermofisher, Cat # A1647801). P2 passage NSCs were plated on poly-L-ornithine (20ug/ml) (Sigma, Cat # P4957) and laminin (5 ug/ml) (Thermo Fisher, Cat # 23017015) coated plates. For 14 days, the cells were differentiated into cortical neurons using neural differentiation medium (NDM) consisting of Neurobasal™ medium, 2 mM L-Glutamine, 1% B-27, 200 µM L-Ascorbic acid (Sigma, Cat # A92902), 0.5 mM c-AMP (Stem Cell Technologies Cat # 73886), 20 ng/ml BDNF (Stem Cell Technologies Cat # 78005), 20 ng/ml GDNF (Stem Cell Technologies Cat # 78058), 20 ng/ml NT-3 (Stem Cell Technologies Cat # 78074), and Penicillin/Streptomycin cocktail. The culture medium was exchanged with fresh NDM every 2-3 days.

### Statistical Analysis

Graphs were generated and statistical analyses performed using Prism 8.4.2 software. All experiments were performed at least three independent times, and statistical analyses were conducted on the composite data unless reported otherwise. Unless otherwise specified, the data were analyzed using a two-way analysis of variance (ANOVA) with Dunnett’s multiple comparisons test to the control (WT_III_). For *in vivo* infections at a single timepoint a one-way ANOVA with Dunnett’s multiple comparisons test to the control (BL6) was used.

---

**Fig. S1.**
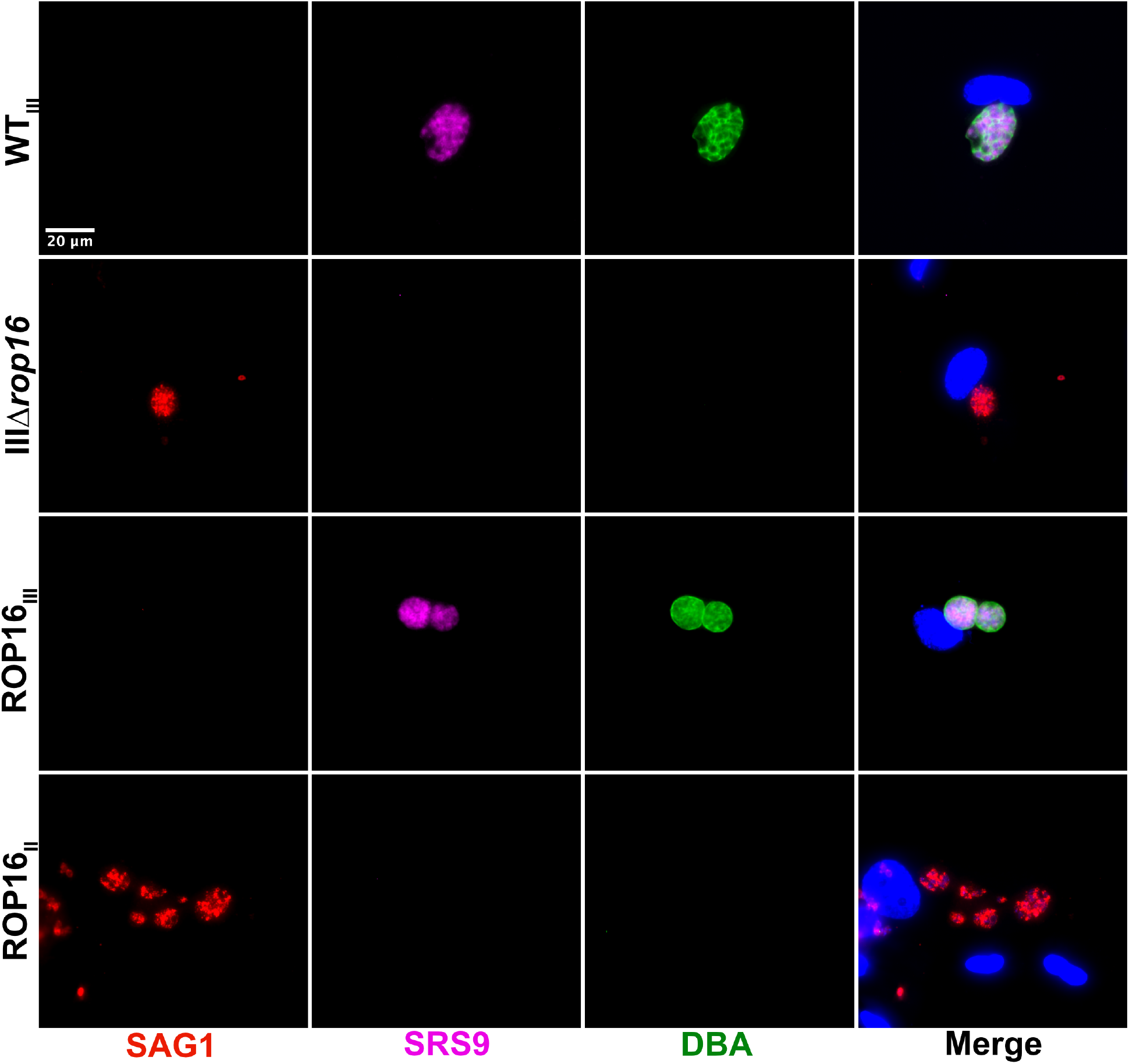
ROP16_III_-deficient parasites are impaired in cyst development in alkaline stressed HFFs. IFA of cyst assay in HFFs. HFFs were infected with the indicated strains for 6 days under alkaline stress. Images depict DBA (green), anti-SAG1 (red), anti-SRS9 (magenta), and DAPI (blue).

**Fig. S2.**
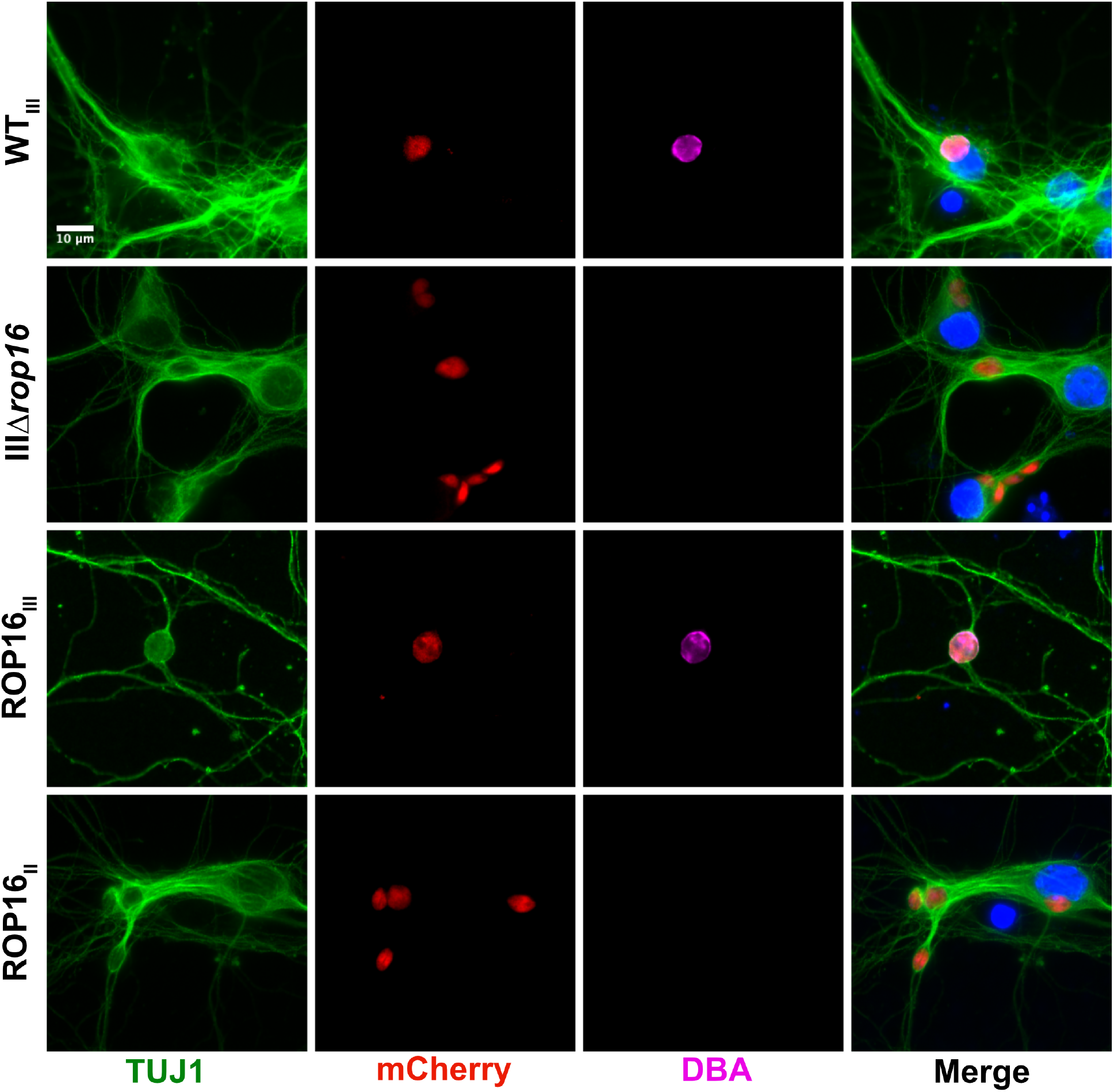
ROP16_III_-deficient parasites are impaired in cyst development in PNCs. IFA of cyst assay in PNCs. PNCs were infected with the indicated strains for 2 days. Images depict anti-TUJ1 (green), mCherry (red), DBA (magenta), and DAPI (blue).

**Fig. S3.**
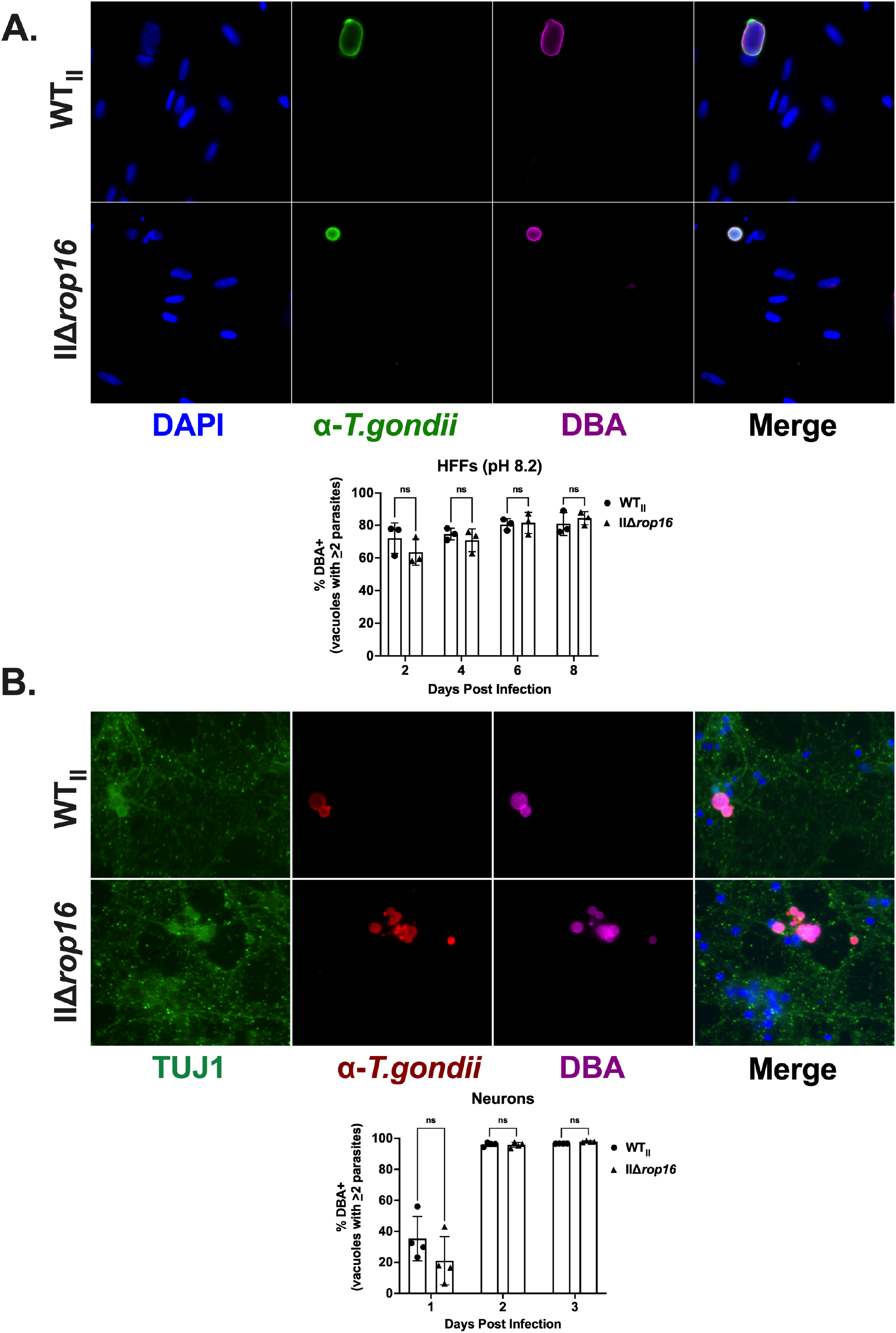
ROP16 is dispensable for cyst formation in type II parasites. (A) Top, IFA of cyst assay in HFFs. HFFs were infected with the indicated strains for 2 days. Images depict anti-*T. gondii* staining (green), DBA (magenta), and DAPI (blue). Bottom, quantification of encystment over time in stress model of encystment. % Cyst as in Fig 1. N = 10 wells/experiment, 3 experiments total. (B) Top, IFA of cyst assay in PNCs. PNCs were infected with the indicated strains for 2 days. Images depict anti-TUJ1 (green), anti-*T. gondii* (red), DBA (magenta), and DAPI (blue). Bottom, quantification of encystment overtime as in (A) except in PNCs. N = 5 wells/experiment, 4 experiments total. Graphs in (A, B): Bars, mean ± SEM. Black dots = Average % cyst for 1 experiment. (A, B) ns = not significant, two-way ANOVA, Dunnett’s multiple comparisons test compared to WT_II_.

**Fig. S4.**
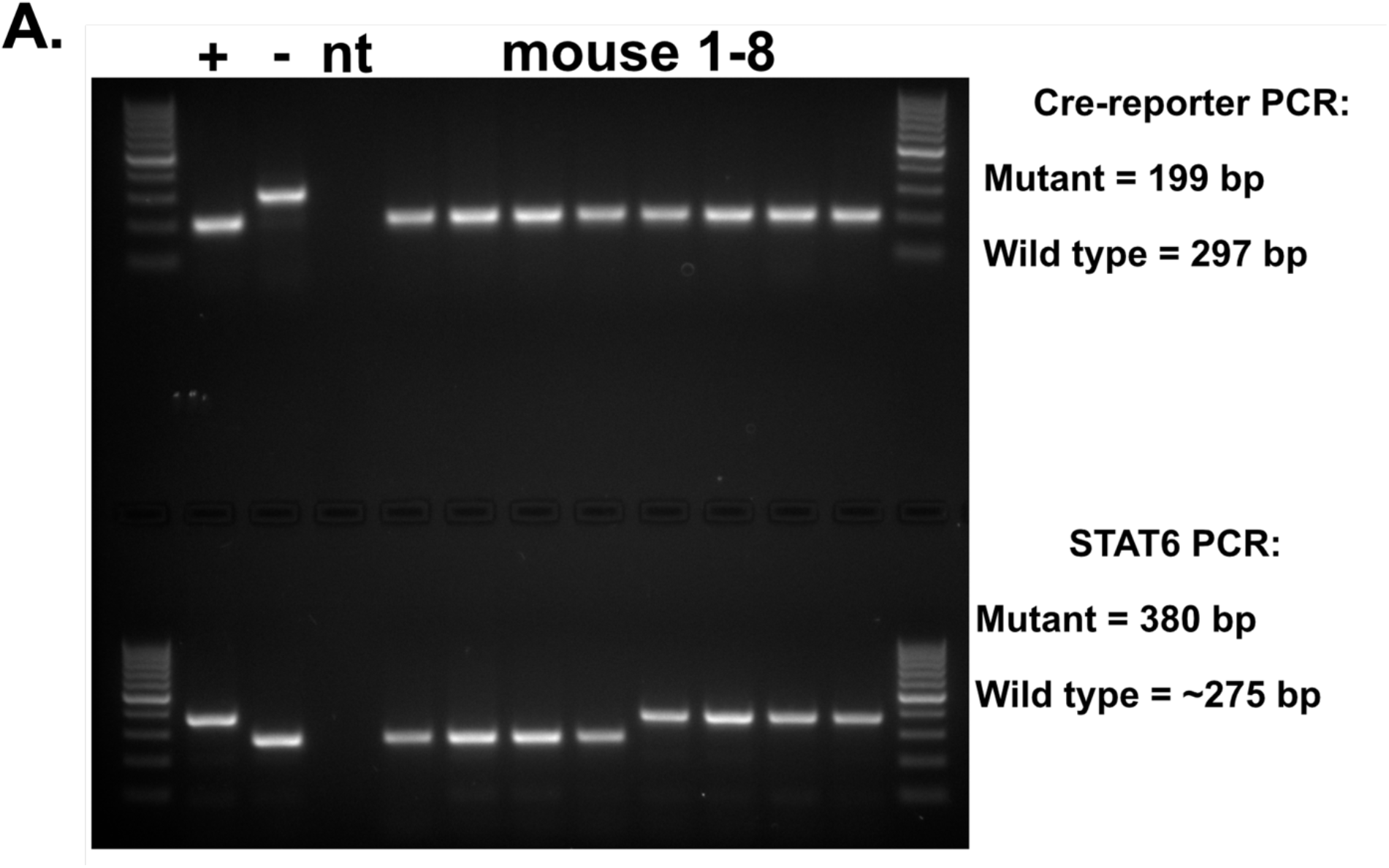
PCR confirmation of STAT6KO Cre reporter mice. Top, PCR for Cre reporter. Wild-type size = 297bp and mutant (Cre reporter) size = 199bp. Bottom, PCR for STAT6. Wild-type (STAT6) size = 275bp and mutant (STAT6KO) size = 380bp. + = positive control, - = negative control and nt = H_2_O control.

